# Efficient management and analysis of large-scale genome-wide data with two R packages: bigstatsr and bigsnpr

**DOI:** 10.1101/190926

**Authors:** Florian Privé, Hugues Aschard, Michael G.B. Blum

## Abstract

**Motivation:** Genome-wide datasets produced for association studies have dramatically increased in size over the past few years, with modern datasets commonly including millions of variants measured in dozens of thousands of individuals. This increase in data size is a major challenge severely slowing down genomic analyses. Specialized software for every part of the analysis pipeline have been developed to handle large genomic data. However, combining all these software into a single data analysis pipeline might be technically difficult.

**Results:** Here we present two R packages, bigstatsr and bigsnpr, allowing for management and analysis of large scale genomic data to be performed within a single comprehensive framework. To address large data size, the packages use memory-mapping for accessing data matrices stored on disk instead of in RAM. To perform data pre-processing and data analysis, the packages integrate most of the tools that are commonly used, either through transparent system calls to existing software, or through updated or improved implementation of existing methods. In particular, the packages implement a fast derivation of Principal Component Analysis, functions to remove SNPs in Linkage Disequilibrium, and algorithms to learn Polygenic Risk Scores on millions of SNPs. We illustrate applications of the two R packages by analysing a case-control genomic dataset for the celiac disease, performing an association study and computing Polygenic Risk Scores. Finally, we demonstrate the scalability of the R packages by analyzing a simulated genome-wide dataset including 500,000 individuals and 1 million markers on a single desktop computer.

**Availability:** https://privefl.github.io/bigstatsr/ & https://privefl.github.io/bigsnpr/

**Contact:** &

**Supplementary information:** Supplementary data are available at *Bioinformatics* online.

## 1 Introduction

Genome-wide datasets produced for association studies have dramatically increased in size over the past years. A range of software and data formats have been developed to perform essential pre-processing steps and data analysis, often optimizing each of these steps within a dedicated implementation. This diverse and extremely rich software environment has been of tremendous benefit for the genetic community. However, it has two limitations: analysis pipelines are becoming very complex and researchers have limited access to diverse analysis tools due to growing data sizes.

Consider first the basic tools necessary to perform a standard genome-wide analysis. Conversions between standard file formats has become a field by itself with several tools such as VCFtools, BCFtools and PLINK, available either independently or incorporated within large framework (Danecek *et al*. 2011; Li *et al*. 2011; Purcell *et al*. 2007). Similarly, quality control software for genome-wide analysis have been developed such as PLINK and the Bioconductor package GWASTools (Gogarten *et al*. 2012). There are also several software for the computation of principal components (PCs) of genotypes, commonly performed to account for population stratification in association studies, including EIGENSOFT (SmartPCA and FastPCA) and FlashPCA (Abraham and Inouye 2014; Abraham *et al*. 2016; Galinsky *et al*. 2016; Price *et al*. 2006). Then, implementation of GWAS analyses also depends on the data format and model analyzed. For example, the software ProbABEL (Aulchenko *et al*. 2010) or SNPTEST (Marchini and Howie 2010) can handle dosage data, while PLINK version 1 is limited to best guess genotypes because of its input file format. Finally, there exists a range of tools for Polygenic Risk Scores (PRSs) such as LDpred (Vilhjálmsson *et al*. 2015) and PRSice (Euesden *et al*. 2015), which provide prediction for quantitative traits or disease risks based on multiple genetic variants. As a result, one has to make extensive bash/perl/R/python scripts to link these software together and convert between multiple file formats, involving many file manipulations and conversions. Overall, this might be a brake on data exploration.

Secondly, increasing size of genetic datasets is the source of major computational challenges and many analytical tools would be restricted by the amount of memory (RAM) available on computers. This is particularly a burden for commonly used analysis languages such as R, Python and Perl. Solving the memory issues for these languages would give access to a broad range of tools for data analysis that have been already implemented. Hopefully, strategies have been developed to avoid loading large datasets in RAM. For storing and accessing matrices, memory-mapping is very attractive because it is seamless and usually much faster to use than direct read or write operations. Storing large matrices on disk and accessing them via memory-mapping has been available for several years in R through “big.matrix” objects implemented in the R package bigmemory (Kane *et al*. 2013). We provide a similar format as filebacked “big.matrix” objects that we called “Filebacked Big Matrices (FBMs)”. Thanks to this matrix-like format, algorithms in R/C++ can be developed or adapted for large genotype data.

## 2 Approach

We developed two R packages, bigstatsr and bigsnpr, that integrate the most efficient algorithms for the pre-processing and analysis of large-scale genomic data while using memory-mapping. Package bigstatsr implements many statistical tools for several types of FBMs (unsigned char, unsigned short, integer and double). This includes implementation of multivariate sparse linear models, Principal Component Analysis, matrix operations, and numerical summaries. The statistical tools developed in bigstatsr can be used for other types of data as long as they can be represented as matrices. Package bigsnpr depends on bigstatsr, using a special type of FBM object to store the genotypes, called “FBM.code256”. Package bigsnpr implements algorithms which are specific to the analysis of SNP arrays, such as calls to external software for processing steps, I/O (Input/Output) operations from binary PLINK files, and data analysis operations on SNP data (thinning, testing, plotting). We use both a real case-control genomic dataset for Celiac disease and large-scale simulated data to illustrate application of the two R packages, including association study and computation of Polygenic Risk Scores. We also compare results from the two R packages with those obtained when using PLINK and EIGENSOFT, and report execution times along with the code to perform major computational tasks.

## 3 Methods

### 3.1 Memory-mapped files

The two R packages don't use standard read operations on a file nor load the genotype matrix entirely in memory. They use an hybrid solution: memory-mapping. Memory-mapping is used to access data, possibly stored on disk, as if it were in memory. This solution is made available within R through the BH package, providing access to Boost C++ Header Files^1^.

We are aware of the software library SNPFile that uses memory-mapped files to store and efficiently access genotype data, coded in C++ (Nielsen and Mailund 2008) and of the R package BEDMatrix^2^ which provides memory-mapping directly for binary PLINK files. With the two packages we developed, we made this solution available in R and in C++ via package Rcpp (Eddelbuettel and François 2011). The major advantage of manipulating genotype data within R, almost as it were a standard matrix in memory, is the possibility of using most of the other tools that have been developed in R (R Core Team 2017). For example, we provide sparse multivariate linear models and an efficient algorithm for Principal Component Analysis (PCA) based on adaptations from R packages biglasso and RSpectra (Qiu and Mei 2016; Zeng and Breheny 2017).

Memory-mapping provides transparent and faster access than standard read/write operations. When an element is needed, a small chunk of the genotype matrix, containing this element, is accessed in memory. When the system needs more memory, some chunks of the matrix are freed from the memory in order to make space for others. All this is managed by the Operating System so that it is seamless and efficient. It means that if the same chunks of data are used repeatedly, it will be very fast the second time they are accessed, the third time and so on. Of course, if the memory size of the computer is larger than the size of the dataset, the file could fit entirely in memory and every second access would be fast.

### 3.2 Data management, preprocessing and imputation

Package bigsnpr currently takes as input a variety of formats (e.g. vcf, bed/bim/fam, ped/map). However, it uses PLINK for conversion to bed/bim/fam format and for Quality Control (QC) of the data, so that we first provide R functions that use system calls to PLINK for the conversion and QC steps (Figure 1). Then, fast read/write operations from/to bed/bim/fam PLINK files are implemented. PLINK files are then read into a “bigSNP” object, which contains the genotype Filebacked Big Matrix (FBM), a data frame with information on samples and another data frame with information on SNPs. We also provide another function which could be used to read from tabular-like text files in order to create a genotype in the format “FBM”.

**Figure 1:**
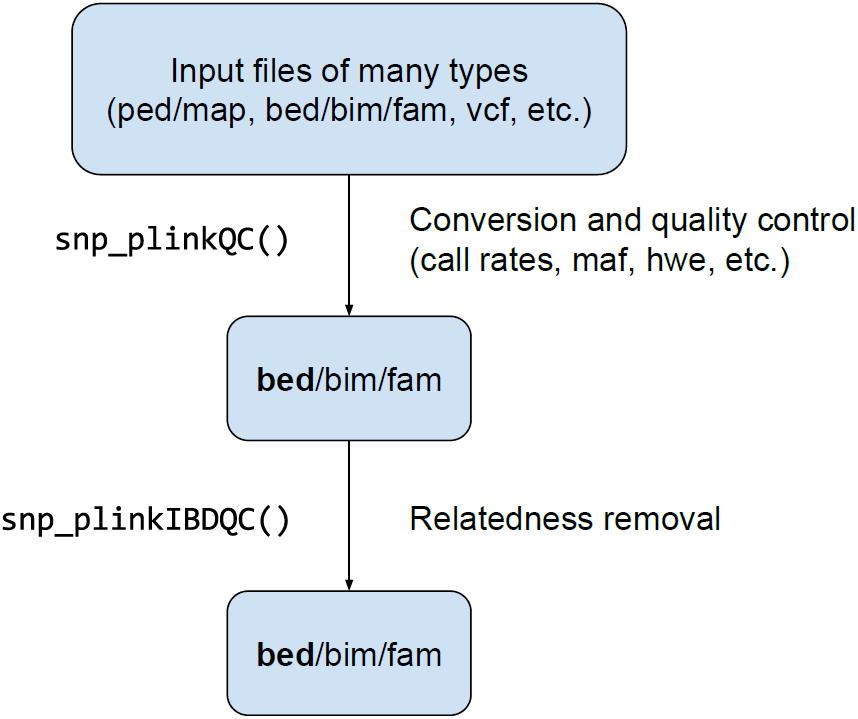
Conversion and Quality Control preprocessing functions available in package bigsnpr via system calls to PLINK.

We developed a special FBM object, called “FBM.code256”, that can be used to seamlessly store up to 256 arbitrary different values, while having a relatively efficient storage. Indeed, each element is stored on one byte which requires 8 times less disk storage than double-precision numbers but 4 times more space than the binary PLINK format “.bed”. With these 256 values, the matrix can store genotype calls and missing values (4 values), best guess genotypes (3 values) and genotype dosages (likelihoods) rounded to two decimal places (201 values).

We also provide two functions for imputing missing values of genotyped SNPs (Figure 2). The first function is a wrapper to PLINK and Beagle (Browning and Browning 2008) which takes bed files as input and return bed files without missing values, and should therefore be used before reading the data in R. The second function is a new algorithm we developed in order to have a fast imputation method without losing much of imputation accuracy. This algorithm is based on Machine Learning approaches for genetic imputation (Wang *et al*. 2012) and doesn’t use phasing, thus allowing for a dramatic decrease in computation time. It only relies on some local XGBoost models. XGBoost is an optimized distributed gradient boosting library that can be used in R and provides some of the best results in machine learning competitions (Chen and Guestrin 2016). XGBoost builds decision trees that can detect nonlinear interactions, partially reconstructing phase, making it well suited for imputing genotype matrices. Systematically, for each SNP, we provide an estimation of imputation error by separating non-missing data into training/test sets. The training set is used to build a model for predicting missing data. The prediction model is then evaluated on the test set for which we know the true genotype values, which gives an unbiased estimator of the number of genotypes that have been wrongly imputed for that particular SNP.

**Figure 2:**
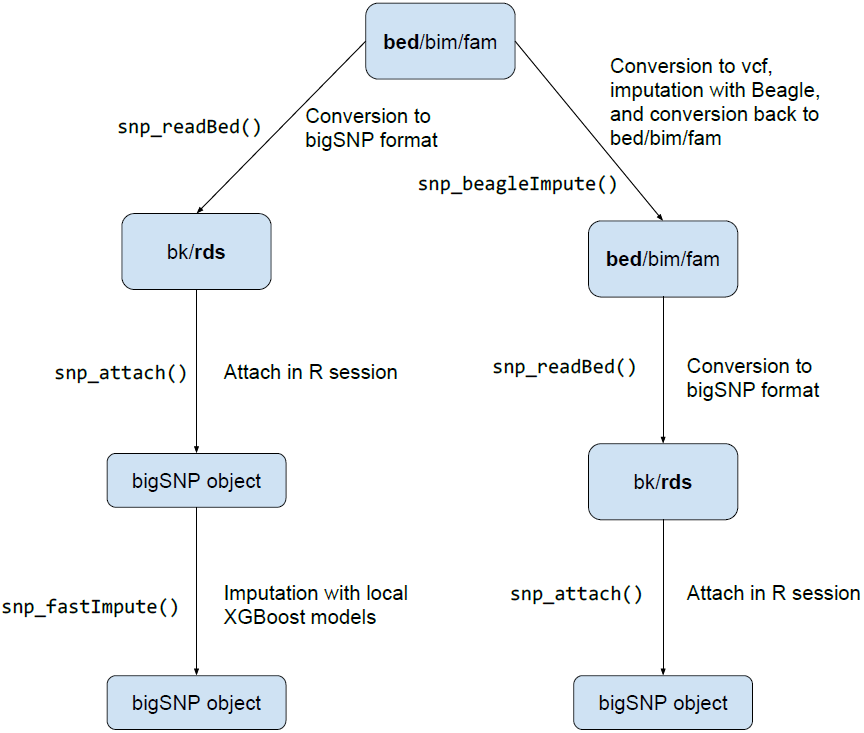
Imputation and reading functions available in package bigsnpr.

### 3.3 Population structure and SNP thinning based on Linkage Disequilibrium

For computing Principal Components (PCs) of a large-scale genotype matrix, we provide several functions related to SNP thinning and two functions, for computing a partial Singular Value Decomposition (SVD), one based on eigenvalue decomposition, big_SVD, and the other on randomized projections, big_randomSVD (Figure 3). While the function based on eigenvalue decomposition is at least quadratic in the smallest dimension, the function based on randomized projections runs in linear time in all dimensions (Lehoucq and Sorensen 1996). Package bigstatsr use the same PCA algorithm as FlashPCA2 called Implicitly Restarted Arnoldi Method (IRAM), which is implemented in R package RSpectra. The main difference between the two implementations is that FlashPCA2 computes vector-matrix multiplications with the genotype matrix based on the binary PLINK file whereas bigstatsr computes these multiplications based on the FBM format, which enables parallel computations and easier subsetting.

**Figure 3:**
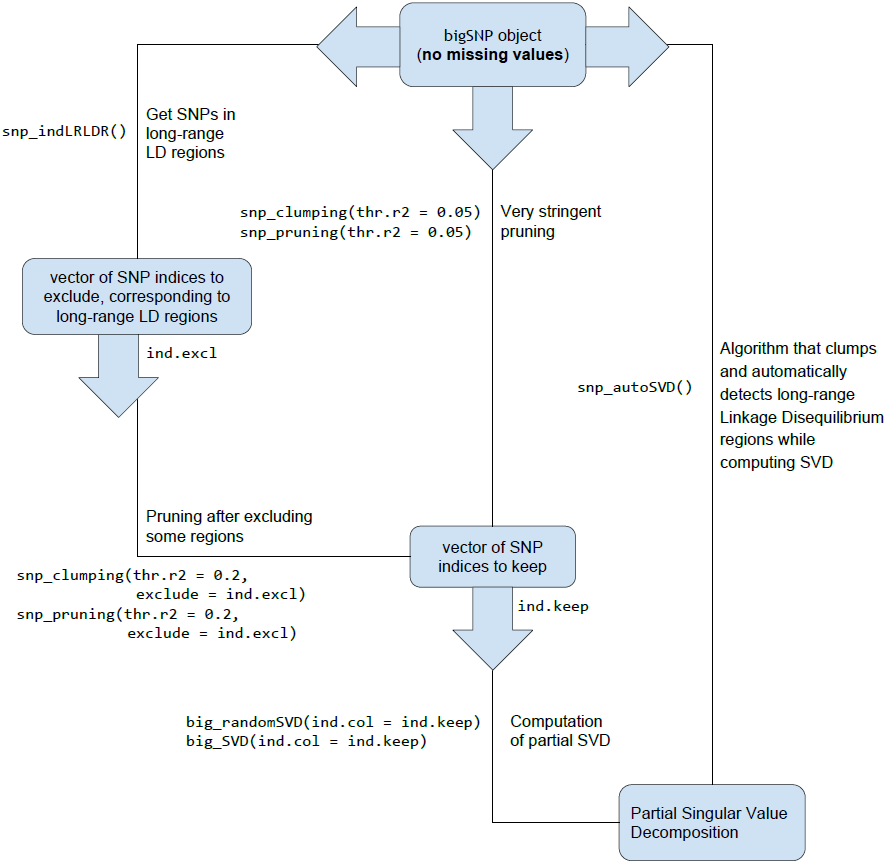
Functions available in packages bigstatsr and bigsnpr for the computation of a partial Singular Value Decomposition of a genotype array, with 3 different methods for thinning SNPs.

SNP thinning improves ascertainment of population structure with PCA (Abdellaoui *et al*. 2013). There are at least 3 different approaches to thin SNPs based on Linkage Disequilibrium, two of them named pruning and clumping, address SNPs in LD close to each others because of recombination events, while the third one address long-range regions with a complex LD pattern due to other biological events such as inversions (Price *et al*. 2008). First, pruning, the most naive approach, is an algorithm that sequentially scan the genome for nearby SNPs in LD, performing pairwise thinning based on a given threshold of correlation. A variant of pruning is clumping. Clumping is useful if a statistic is available to sort the SNPs by importance, e.g. association with a phenotype, and for discarding SNPs in LD with a more associated SNP relatively to the phenotype of interest. Furthermore, we advise to always use clumping instead of pruning (by using the minor allele frequency as the statistic of importance, which is the default) because, in some particular cases, pruning can leave regions of the genome without any representative SNP at all^3^.

As mentioned above, the third approach that is generally combined with pruning or clumping consists of removing SNPs in long-range LD regions (Price *et al*. 2008). Long-range LD regions for the human genome are available as an online table that our packages can use to discard SNPs in long-range LD regions while computing PCs^4^. However, the pattern of LD might be population specific, so we developed an algorithm that automatically detects these regions and removes them. This algorithm consists in the following steps: first, PCA is performed using a subset of SNP remaining after clumping, then outliers SNPs are detected using Mahalanobis distance as implemented in the R package pcadapt (Luu *et al*. 2017). Finally, the algorithm considers that consecutive outlier SNPs are in long-range LD regions. Indeed, a long-range LD region would cause SNPs in this region to have strong consecutive weights (loadings) in the PCA. This algorithm is implemented in function snp_autoSVD and will be referred by this name in the rest of the paper.

### 3.4 Association tests and Polygenic Risk Scores

Any test statistic that is based on counts could be easily implemented because we provide fast counting summaries. Among these tests, the Armitage trend test and the MAX3 test statistic are already provided for binary outcome (Zheng *et al*. 2012). We also implement statistical tests based on linear and logistic regressions. For the linear regression, for each SNP *j*, a t-test is performed on *β*^(*j*)^ where

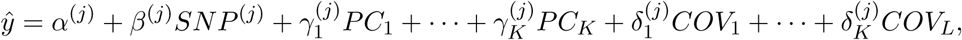

and *K* is the number of principal components and *L* is the number of other covariates (such as the age and gender). Similarly, for the logistic regression, for each SNP *j*, a Z-test is performed on *β*^(*j*)^ where

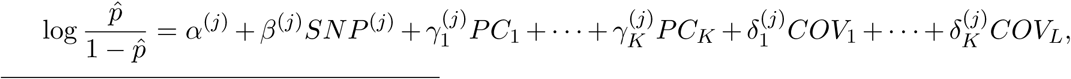

and *p̂* = ℙ(*Y* = 1) and *Y* denotes the binary phenotype.

The R packages also implement functions to compute Polygenic Risk Scores using two approaches. The first method is the widely-used Pruning + Thresholding (P+T) model based on univariate GWAS summary statistics as described in previous equations. Under the P+T model, a coefficient of regression is learned independently for each SNP along with a corresponding p-value. The SNPs are first clumped (P) so that there remains only SNPs that are weakly correlated with each other. Thresholding (T) consists in removing SNPs that are under a certain level of significance (P-value threshold to be determined). A polygenic risk score is defined as the sum of allele counts of the remaining SNPs weighted by the corresponding regression coefficients (Chatterjee *et al*. 2013; Dudbridge 2013; Golan and Rosset 2014). The second approach doesn’t use univariate summary statistics but instead train a multivariate model on all the SNPs and covariables at once, optimally accounting for correlation between predictors (Abraham *et al*. 2012). The currently available models are linear and logistic regressions and Support Vector Machine (SVM). These models include lasso and elastic-net regularizations, which reduce the number of predictors (SNPs) included in the predictive models (Friedman *et al*. 2010; Tibshirani 1996; Zou and Hastie 2005). Package bigstatsr provides a fast implementation of these models by using efficient rules to discard most of the predictors (Tibshirani *et al*. 2012). The implementation of these algorithms is based on modified versions of functions available in the R packages sparseSVM and biglasso (Zeng and Breheny 2017). These modifications allow to include covariates in the models and to use these algorithms on the special type of FBM called “FBM.code256” used in bigsnpr.

### 3.5 Data analyzed

In this paper, two datasets are analyzed: the celiac disease cohort and the POPRES datasets (Dubois *et al*. 2010; Nelson *et al*. 2008). The Celiac dataset is composed of 15,283 individuals of European ancestry genotyped on 295,453 SNPs. The POPRES dataset is composed of 1385 individuals of European ancestry genotyped on 447,245 SNPs. For computation times comparison, we replicated individuals in the Celiac dataset 5 and 10 times in order to increase sample size while keeping the same population structure and pattern of Linkage Disequilibrium as the original dataset. To assess scalibility of our algorithms for a biobank-scale genotype dataset, we formed another dataset of 500,000 individuals and 1 million SNPs, also through replication of the Celiac dataset.

### 3.6 Reproducibility

All the code used in this paper along with results, such as execution times and figures, are available as HTML R notebooks in the Supplementary Data.

## 4 Results

### 4.1 Overview

We present the results for three different analyses. First, we illustrate the application of R packages bigstatsr and bigsnpr. Secondly, we compare the performance of the R packages to the performance obtained with PLINK and FastPCA (EIGENSOFT). Thirdly, we present results of the two new methods implemented in these packages, one method for the automatic detection and removal of long-range LD regions in PCA and another for the imputation of missing genotypes. We use three types of data: a case-control cohort for the celiac disease, the European population cohort POPRES and simulated datasets using real genotypes from the Celiac cohort. We compare performances on two computers, a desktop computer with 64GB of RAM and 12 cores (6 physical cores), and a laptop with only 8GB of RAM and 4 cores (2 physical cores). For the functions that enable parallelism, we use half of the cores available on the corresponding computer.

### 4.2 Application

We performed an association study and computed a polygenic risk score for the Celiac cohort. The data was preprocessed following steps from figure 1, removing individuals and SNPs which had more than 5% of missing values, non-autosomal SNPs, SNPs with a minor allele frequency lower than 0.05 or a p-value for the Hardy-Weinberg exact test lower than 10^‒10^, and finally, removing the first individual in each pair of individuals with a proportion of alleles shared IBD greater than 0.08 (Purcell *et al*. 2007). For the POPRES dataset, this resulted in 1382 individuals and 344,614 SNPs with no missing value. For the Celiac dataset, this resulted in 15,155 individuals and 281,122 SNPs with an overall genotyping rate of 99.96%. The 0.04% missing genotype values were imputed with the XGBoost method. If we used a standard R matrix to store the genotypes, this data would require 32GB of memory. On the disk, the “.bed” file requires 1GB and the “.bk” file (storing the FBM) requires 4GB.

We used bigstatsr and bigsnpr R functions to compute the first Principal Components (PCs) of the Celiac genotype matrix and to visualize them (Figure 4). We then performed a Genome-Wide Association Study (GWAS) investigating how Single Nucleotide Polymorphisms (SNPs) are associated with the celiac disease, while adjusting for PCs, and plotted the results as a Manhattan plot (Figure 5). As illustrated in the supplementary data, the whole pipeline is user-friendly and requires only 20 lines of R code.

**Figure 4:**
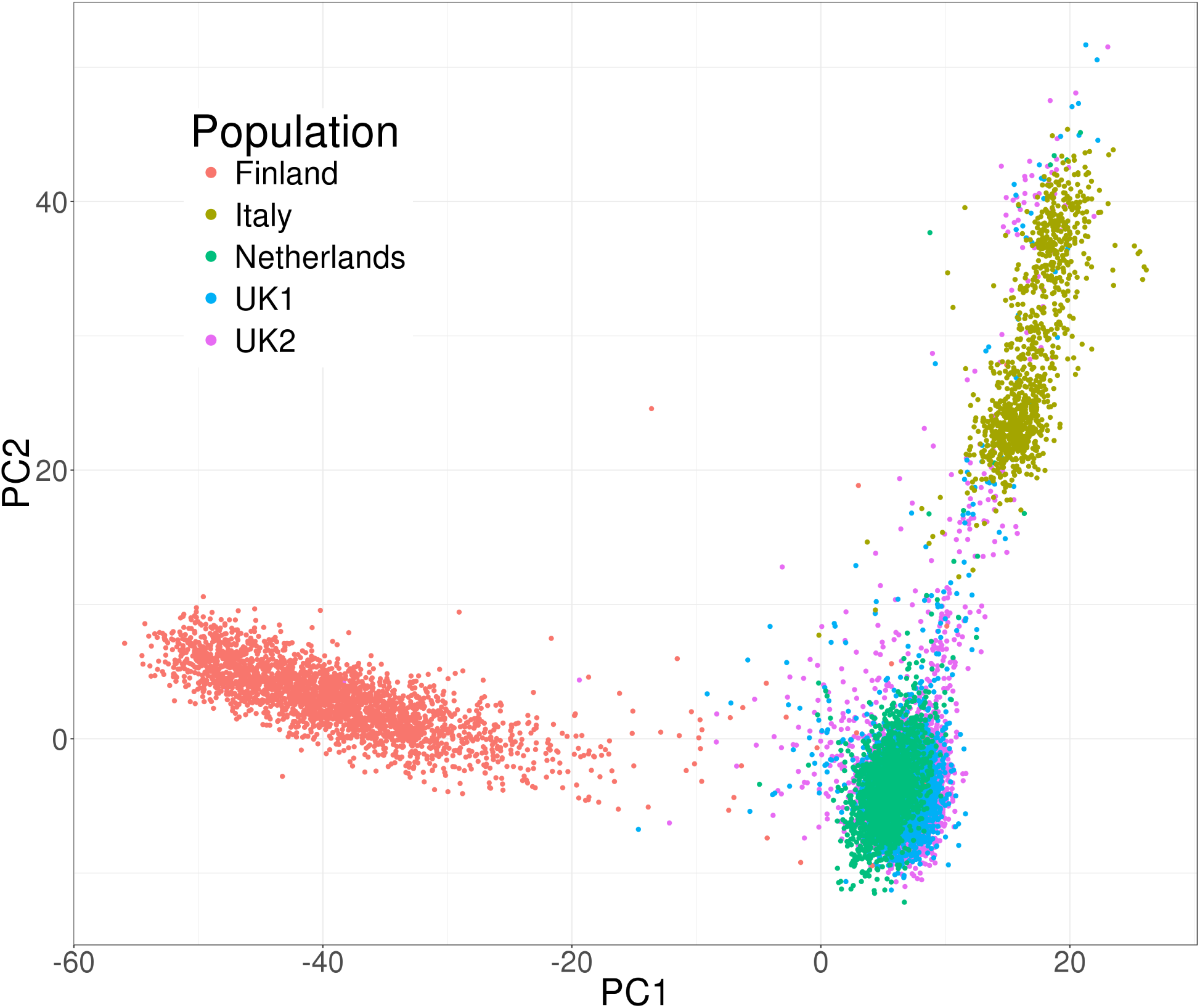
Principal Components of the celiac cohort genotype matrix produced by package bigstatsr.

**Figure 5:**
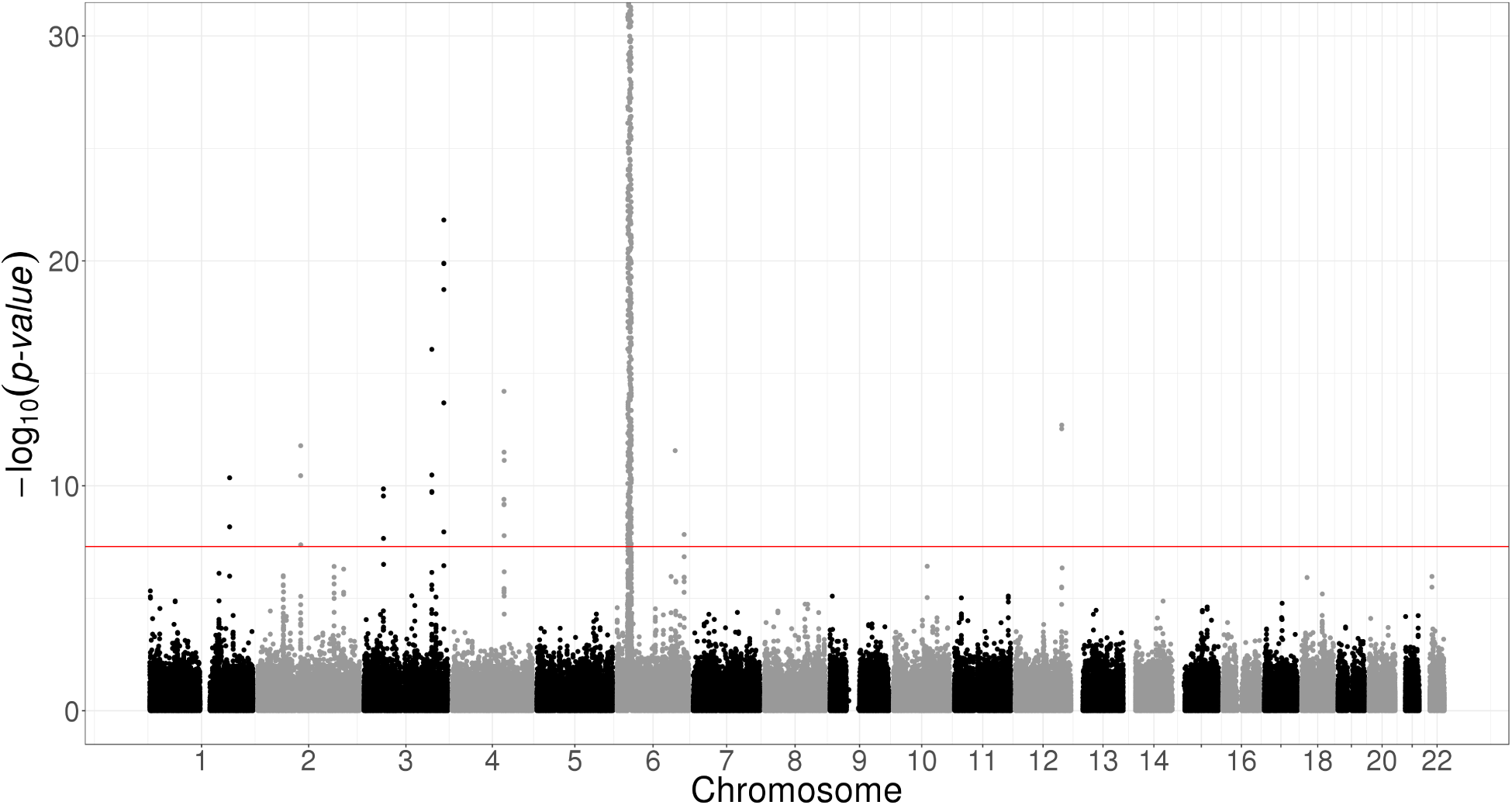
Manhattan plot of the celiac disease cohort produced by package bigsnpr. Some SNPs in chromosome 6 have p-values smaller than the 10^‒30^ threshold used for vizualisation purposes.

To illustrate the scalability of the two R packages, we performed a GWAS analysis on 500K individuals and 1M SNPs. The GWAS analysis completed in approximately 11 hours using the aforementioned desktop computer. The GWAS analysis was composed of four main steps. First we read from PLINK files in our format "bigSNP" in 1 hour. Then, we removed SNPs in long-range LD regions and used SNP clumping, leaving 93,083 SNPs in 5.4h. Then, the 10 first PCs were computed on the 500K individuals and these remaining SNPs in 1.8h. Finally, we performed a linear association test on the complete 500K dataset for each of the 1M SNPs, using the 10 first PCs as covariables in 2.9h.

### 4.3 Method Comparison

We first compared the GWAS and PRS computations obtained with the R packages to the ones obtained with PLINK 1.9 and EIGENSOFT 6.1.4. For most functions, multithreading is not available yet in PLINK, nevertheless, PLINK-specific algorithms that use bitwise parallelism (e.g. pruning) are still faster than the parallel algorithms reimplemented in package bigsnpr (Table 1). Overall, the computations with our two R packages for an association study and a polygenic risk score are of the same order of magnitude as when using PLINK and EIGEN-SOFT (Tables 1 and 2). However, the whole analysis pipeline makes use of R calls only; there is no need to write temporary files and functions have parameters which enable subsetting of the genotype matrix without having to copy it.

**Table 1:** Execution times with bigstatsr and bigsnpr compared to PLINK and FastPCA for making a GWAS for the Celiac dataset. The first execution time is with a desktop computer (6 cores used and 64GB of RAM) and the second one is with a laptop computer (2 cores used and 8GB of RAM).

| Operation | Execution times (in seconds) |  |
| --- | --- | --- |
|  | PLINK and FastPCA | bigstatsr and bigsnpr |
| Reading PLINK files | n/a | 5 / 20 |
| Pruning | 4 / 4 | 14 / 52 |
| Computing 10 PCs | 306 / 315 | 58 / 180 |
| GWAS (binary phenotype) | 339 / 293 | 301 / 861 |
| Total | 649 / 612 | 378 / 1113 |

**Table 2:** Execution times with bigstatsr and bigsnpr compared to PLINK and FastPCA for making a PRS for a training set of 80% of the Celiac dataset. The first execution time is with a desktop computer (6 cores used and 64GB of RAM) and the second one is with a laptop computer (2 cores used and 8GB of RAM).

| Operation | Execution times (in seconds) |  |
| --- | --- | --- |
|  | PLINK | bigstatsr and bigsnpr |
| GWAS (binary phenotype) | 232 / 239 | 178 / 650 |
| Clumping | 49 / 58 | 10 / 35 |
| PRS | 9 / 10 | 2 / 3 |
| Total | 290 / 307 | 190 / 688 |

On our desktop computer, we compared the computation times of FastPCA, FlashPCA2 to the similar function big_randomSVD implemented in bigstatsr. For each comparison, we used the 93,083 SNPs which were remaining after pruning and we computed 10 PCs. We used the datasets of growing size simulated from the Celiac dataset. Overall, our function big_randomSVD showed to be almost twice as fast as FastPCA and FlashPCA2 and 8 times as fast when using parallelism (an option not currently possible with either FastPCA or Flash-PCA2) with 6 cores (Figure 6). We also compared results in terms of precision by comparing squared correlation between approximated PCs and “true” PCs provided by an exact singular value decomposition obtained with SmartPCA. FastPCA, FlashPCA2 and bigstatsr infer the true first 6 PCs but the squared correlation between true PCs and approximated ones decreases for larger PCs when using FastPCA (Fast mode of EIGENSOFT) whereas it remains larger than 0.999 when using FlashPCA2 or bigstatsr (Figure 7).

**Figure 6:**
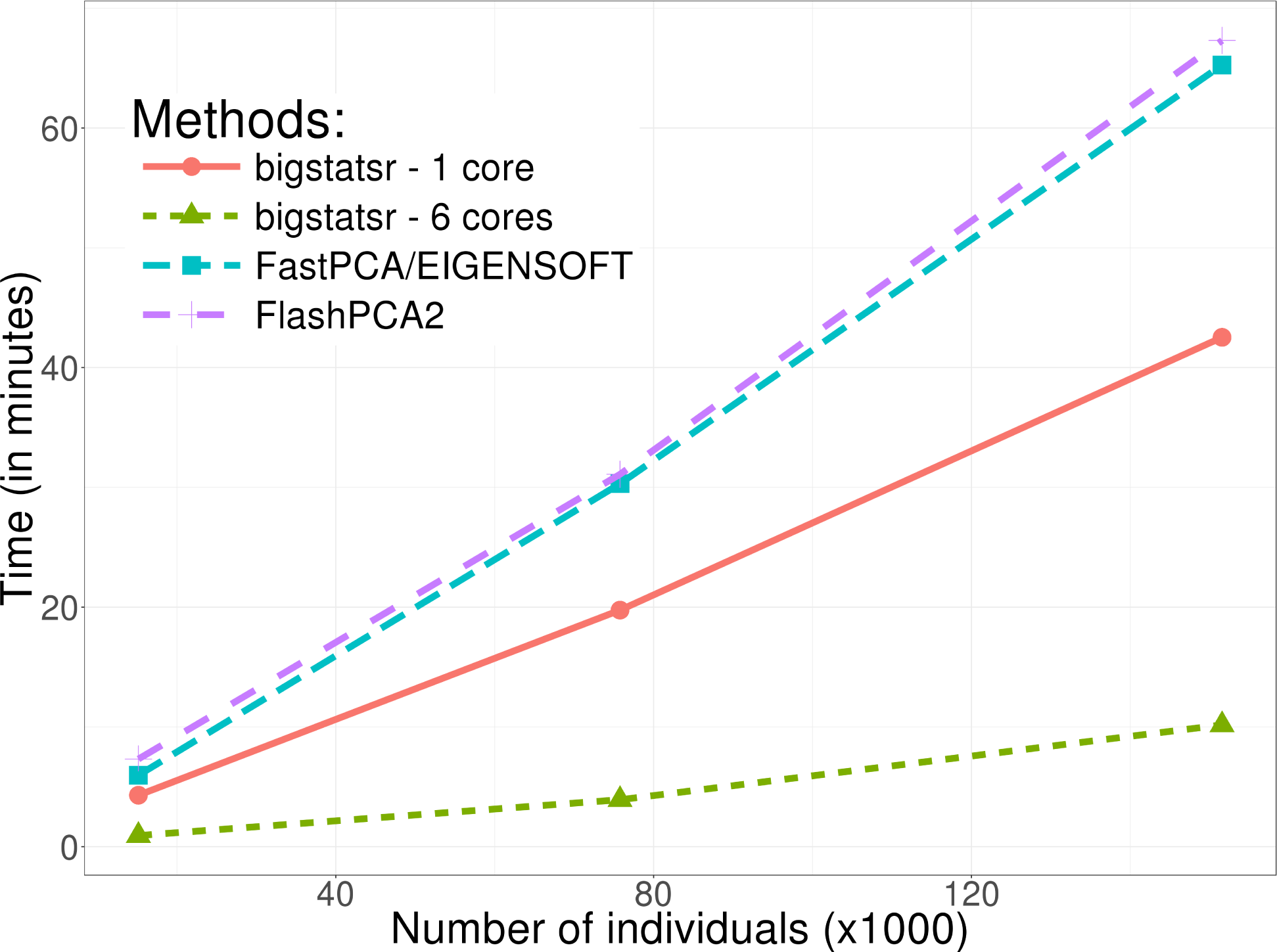
Benchmark comparisons between randomized Partial Singular Value Decomposition available in FlashPCA2, FastPCA (fast mode of SmartPCA/EIGENSOFT) and package bigstatsr. It shows the computation time in minutes as a function of the number of samples. The first 10 principal components have been computed based on the 93,083 SNPs which remained after thinning.

**Figure 7:**
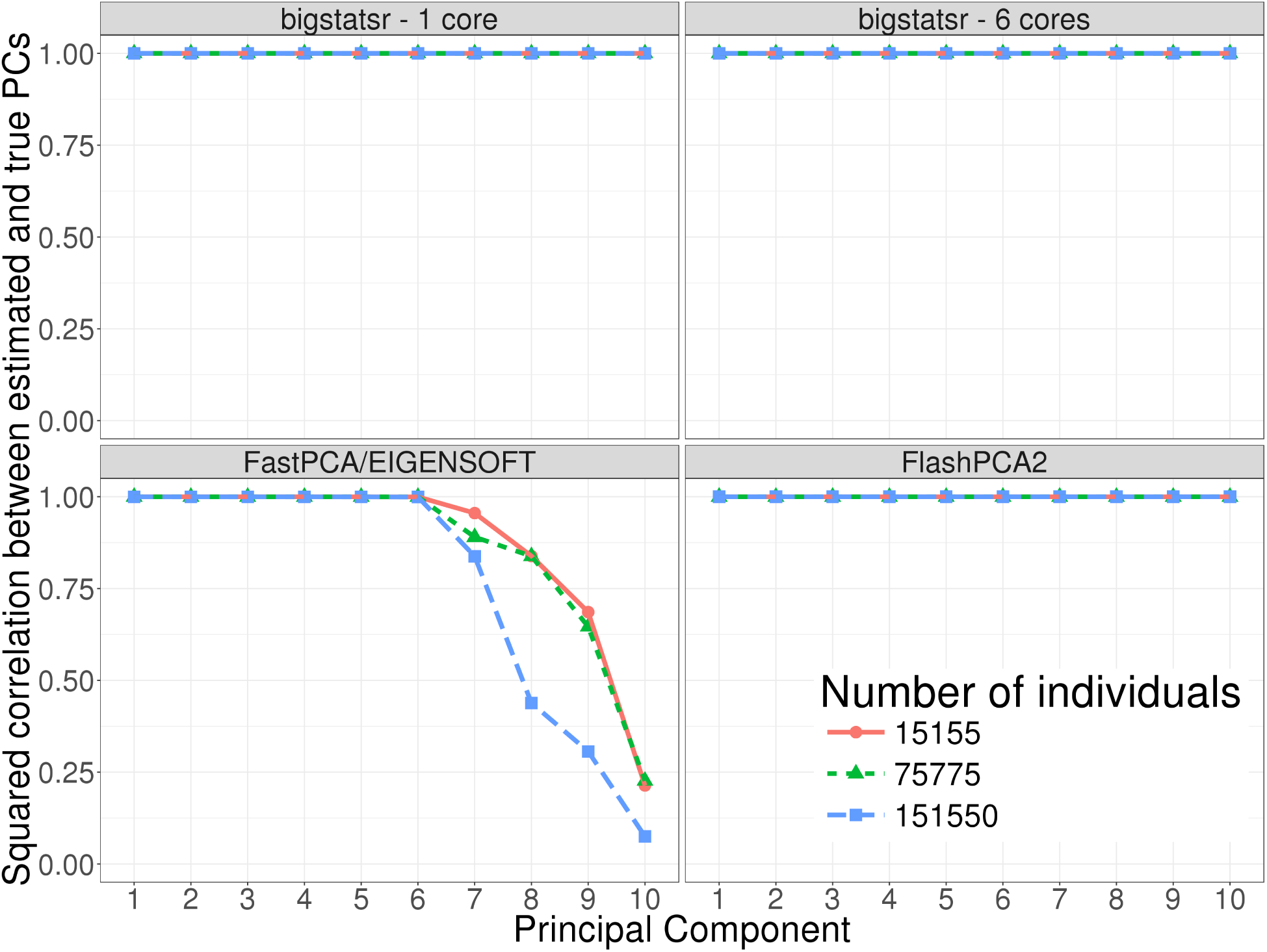
Precision comparisons between randomized Partial Singular Value Decomposition available in FlashPCA2, FastPCA (fast mode of SmartPCA/EIGENSOFT) and package bigstatsr. It shows the squared correlation between approximated PCs and “true” PCs (given by the slow mode of SmartPCA) of the Celiac dataset (whose individuals have been repeated 1, 5 and 10 times).

### 4.4 Automatic detection of long-range LD regions

For the detection of long-range LD regions during the computation of PCA, we tested the function snp_autoSVD on both the Celiac and POPRES datasets. For the POPRES dataset, the algorithm converged in two iterations. The first iterations found 3 long-range LD regions in chromosomes 2, 6 and 8 (Table S1). We compared the PCs of genotypes obtained after applying snp_autoSVD with the PCs obtained after removing pre-determined long-range LD regions^5^ and found a mean correlation of 89.6% between PCs, mainly due to a rotation of PC7 and PC8 (Table S2). For the Celiac dataset, we found 5 long-range LD regions (Table S3) and a mean correlation of 98.6% between PCs obtained with snp_autoSVD and the ones obtained by clumping with removing of predetermined long-range LD regions (Table S4).

For the Celiac dataset, we further compared results of PCA obtained when using snp_autoSVD and when computing PCA without removing any long range LD region (only clumping at *R*^2^ > 0.2). When not removing any long range LD region, we show that PC4 and PC5 don’t capture population structure and correspond to a long-range LD region in chromosome 8 (Figures S1 and S2). When automatically removing some long-range LD regions with snp_autoSVD, we show that PC4 and PC5 reflect population structure (Figure S1). Moreover, loadings are more equally distributed among SNPs after removal of long-range LD regions (Figure S2). This is confirmed by Gini coefficients (measure of dispersion) of each squared loadings that are significantly smaller when computing SVD with snp_autoSVD than when no long-range LD region is removed (Figure S3).

### 4.5 Imputation of missing values for genotyped SNPs

For the imputation method based on XGBoost, we compared the imputation accuracy and computation times with Beagle on the POPRES dataset. The histogram of the minor allele frequencies (MAFs) of this dataset is provided in figure S4 and there is no missing value. We used a Beta-binomial distribution to simulate the number of missing values by SNP and then randomly introduced missing values according to these numbers, resulting in approximately 3% of missing values overall (Figure S5). Imputation was compared between function snp_fastImpute of package bigsnpr and Beagle 4.1 (version of January 21, 2017). Overall, snp_fastImpute made 4.7% of imputation errors whereas Beagle made only 3.1% of errors but it took Beagle 14.6 hours to complete while our method only took 42 minutes (20 times less). We also show that the estimation of the number of imputation errors is accurate (Figure S6). For the Celiac dataset in which there was already missing values, in order to further compare computation times, we report that snp_fastImpute took less than 10 hours to complete for the whole genome whereas Beagle didn’t finish imputing chromosome 1 in 48 hours.

## 5 Discussion

We have developed two R packages, bigstatsr and bigsnpr, which enable multiple analyses of large-scale genotype datasets in a single comprehensive framework. Linkage Disequilibrium pruning, Principal Component Analysis, association tests and computations of polygenic risk scores are made available in this software. Implemented algorithms are both fast and memory-efficient, allowing the use of laptops or desktop computers to make genome-wide analyses. Technically, bigstatsr and bigsnpr could handle any size of datasets. However, if the OS has to often swap between the file and the memory for accessing the data, this would slow down data analysis. For example, the Principal Component Analysis (PCA) algorithm in bigstatsr is iterative so that the matrix has to be sequentially accessed over a hundred times. If the number of samples times the number of SNPs remaining after pruning is larger than the available memory, this slowdown would happen. For instance, a 32GB computer would be slow when computing PCs on more than 100K samples and 300K SNPs remaining after LD thinning.

The two R packages use a matrix-like format, which makes it easy to develop new functions in order to experiment and develop new ideas. Integration in R makes it possible to take advantage of the vast and diverse R libraries. For example, we developed a fast and accurate imputation algorithm for genotyped SNPs using the widely-used machine learning algorithm XGBoost available in the R package xgboost. Other functions, not presented here, are also available and all the functions available within the package bigstatsr are not specific to SNP arrays, so that they could be used for other omic data or in other fields of research.

We think that the two R packages and the corresponding data format could help researchers to develop new ideas and algorithms to analyze genome-wide data. For example, we wish to use these packages to train much more accurate predictive models than the standard P+T model currently in use when computing Polygenic Risk Scores. As a second example, multiple imputation has been shown to be a very promising method for increasing statistical power of a GWAS (Palmer and Pe’er 2016), and it could be implemented with the data format “FBM.code256” without having to write multiple files.

## Acknowledgements

Authors acknowledge Grenoble Alpes Data Institute, supported by the French National Research Agency under the “Investissements d?avenir” program (ANR-15-IDEX-02) and the LabEx PERSYVAL-Lab (ANR-11-LABX-0025-01).

## Supplementary Data

### 5.1 Long-range LD regions

**Table S1:**
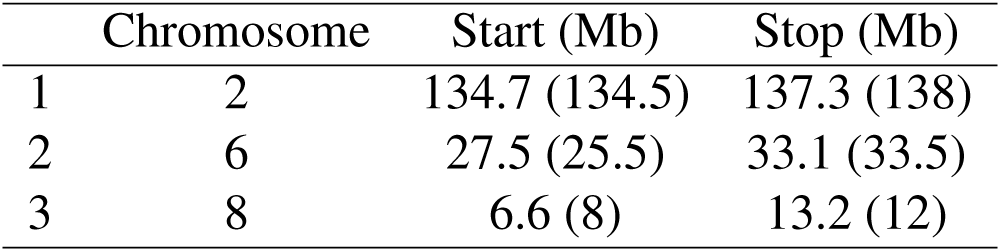
Regions found by snp_autoSVD for the POPRES dataset. Numbers in parentheses correspond to regions referenced in Price *et al*. (2008).

**Table S2:** Correlation between scores of PCA for the POPRES dataset when automatically removing long-range LD regions and when removing them based on a predefined table.

|  | PC1 | PC2 | PC3 | PC4 | PC5 | PC6 | PC7 | PC8 | PC9 | PC10 |
| --- | --- | --- | --- | --- | --- | --- | --- | --- | --- | --- |
| PC1 | 100.0 | -0.1 | -0.0 | 0.1 | -0.1 | 0.0 | 0.0 | 0.0 | -0.0 | -0.0 |
| PC2 | 0.1 | 100.0 | -0.0 | 0.1 | -0.0 | -0.0 | -0.0 | 0.2 | -0.1 | -0.0 |
| PC3 | 0.0 | -0.0 | 99.9 | 0.9 | 0.1 | -0.1 | -0.3 | 0.2 | 0.4 | 0.1 |
| PC4 | -0.1 | -0.1 | -0.9 | 99.7 | -1.0 | 0.7 | 0.6 | 0.2 | 0.3 | 0.9 |
| PC5 | 0.1 | 0.0 | -0.1 | 1.1 | 99.3 | 1.3 | -0.8 | 1.3 | -4.2 | -2.4 |
| PC6 | -0.0 | 0.0 | 0.1 | -0.7 | -1.0 | 97.7 | -3.5 | 6.1 | 7.9 | -6.2 |
| PC7 | -0.0 | -0.1 | 0.2 | -0.3 | -1.7 | 0.3 | 58.3 | 73.2 | -25.9 | 9.1 |
| PC8 | 0.1 | -0.1 | -0.3 | 0.4 | -0.5 | -5.3 | -73.5 | 59.5 | 15.8 | 13.2 |
| PC9 | 0.0 | 0.1 | -0.4 | -0.8 | 5.0 | -7.6 | 27.8 | 11.0 | 91.9 | 9.0 |
| PC10 | 0.1 | 0.0 | 0.0 | -0.9 | 1.6 | 10.2 | 3.9 | -19.6 | -6.3 | 89.2 |

**Table S3:**
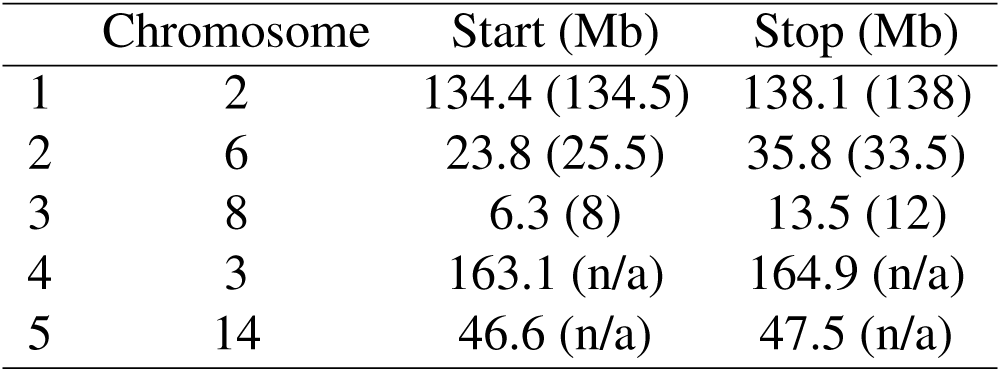
Regions found by snp_autoSVD for the celiac dataset. Numbers in parentheses correspond to regions referenced in Price *et al*. (2008).

**Table S4:** Correlation between scores of PCA for the Celiac dataset when automatically removing long-range LD regions and when removing them based on a predefined table.

|  | PC1 | PC2 | PC3 | PC4 | PC5 | PC6 | PC7 | PC8 | PC9 | PC10 |
| --- | --- | --- | --- | --- | --- | --- | --- | --- | --- | --- |
| PC1 | 100.0 | -0.1 | -0.1 | 0.0 | 0.0 | 0.0 | 0.0 | 0.0 | 0.0 | 0.0 |
| PC2 | 0.1 | 100.0 | 0.0 | 0.0 | -0.0 | -0.0 | 0.0 | -0.0 | -0.0 | -0.0 |
| PC3 | 0.1 | -0.0 | 99.9 | 0.2 | -0.0 | 0.1 | 0.1 | 0.1 | 0.0 | -0.1 |
| PC4 | -0.0 | -0.0 | -0.3 | 99.9 | -0.1 | 0.1 | -0.1 | 0.0 | 0.1 | 0.1 |
| PC5 | 0.0 | 0.0 | 0.0 | 0.1 | 99.7 | 0.9 | -0.3 | 0.1 | -0.8 | -0.6 |
| PC6 | -0.0 | 0.0 | -0.1 | -0.2 | -0.8 | 99.6 | 0.5 | -0.5 | -0.2 | -0.4 |
| PC7 | -0.0 | 0.0 | -0.1 | 0.0 | 0.5 | -0.4 | 98.9 | 3.1 | 0.7 | 1.6 |
| PC8 | 0.0 | 0.0 | -0.2 | -0.0 | -0.2 | 0.5 | -3.2 | 98.4 | -4.5 | -1.5 |
| PC9 | -0.0 | -0.0 | -0.0 | 0.0 | 0.6 | 0.1 | -0.7 | 4.6 | 96.9 | -10.7 |
| PC10 | -0.0 | -0.0 | 0.1 | -0.1 | 0.3 | 0.1 | -1.2 | 1.5 | 8.6 | 92.7 |

**Figure S1:**
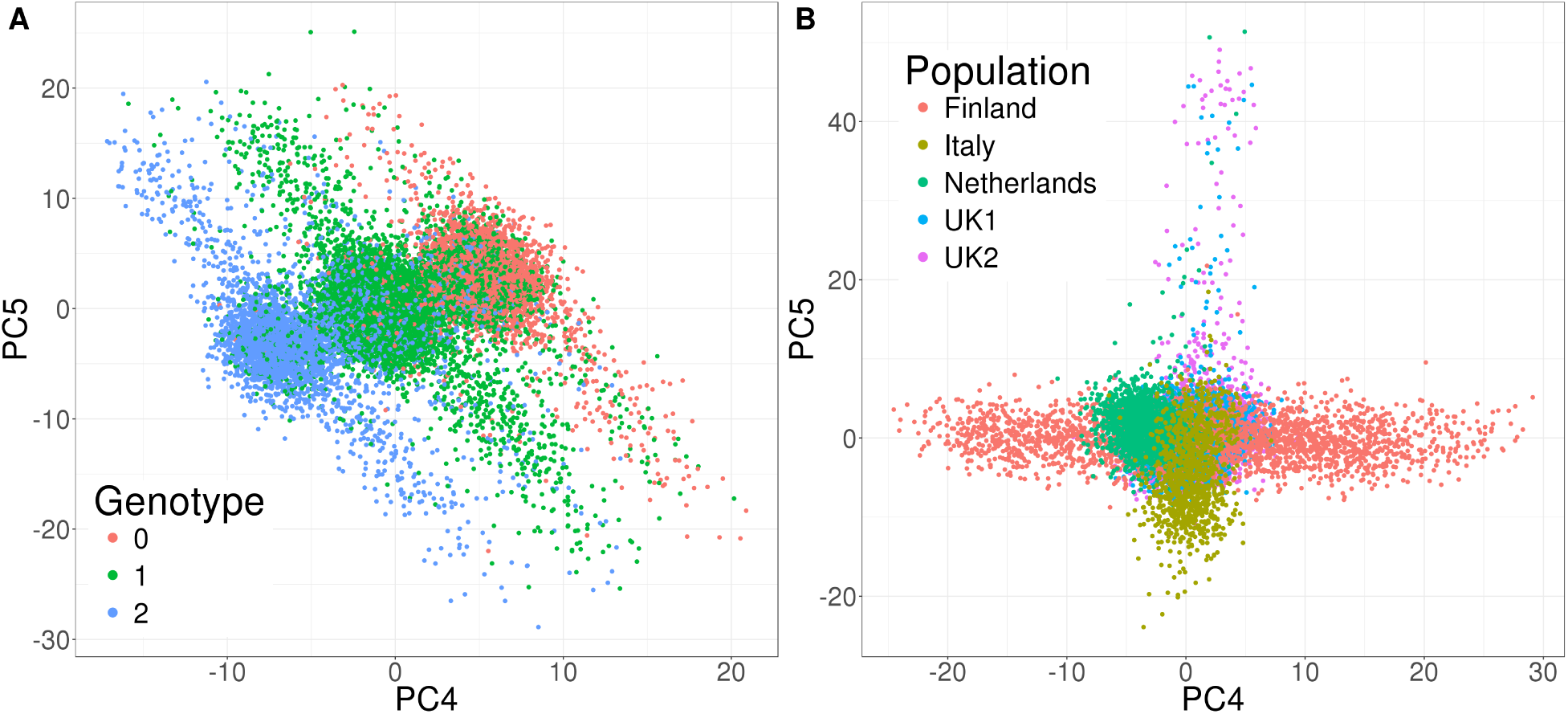
PC4 and PC5 of the celiac disease dataset. Left panel, PC scores obtained without removing any long range LD region (only clumping at *R*^2^ > 0.2). Individuals are coloured according to their genotype at the SNP that has the highest loading for PC4. Right panel, PC scores obtained with the automatic detection and removal of long-range LD regions. Individuals are coloured according to their population of origin.

### 5.2 Imputation

**Figure S2:**
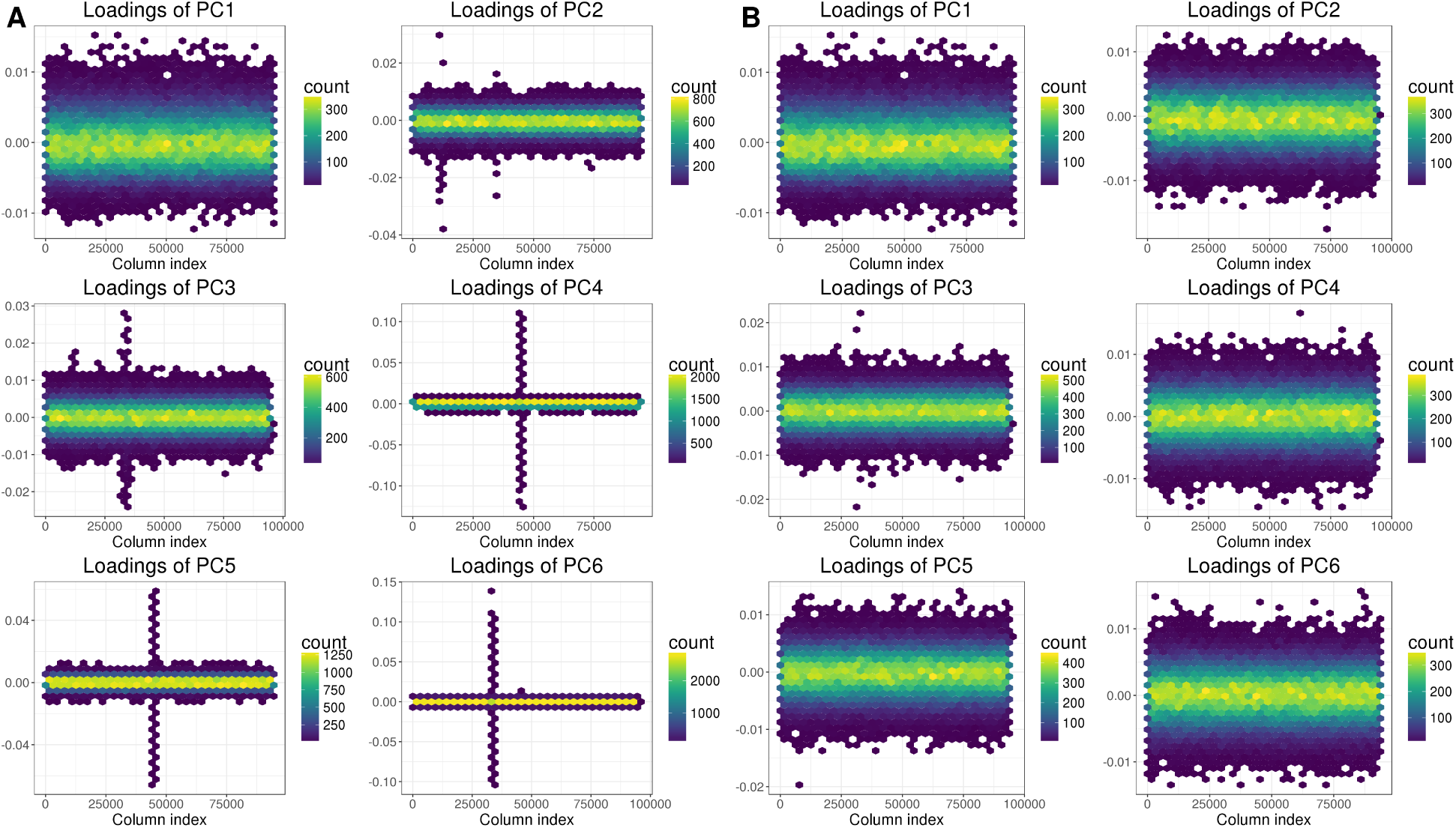
Loadings of first 6 PCs of the celiac disease dataset plotted as hexbins (2-D histogram with hexagonal cells). On the left, without removing any long range LD region (only clumping at *R*^2^ > 0.2). On the right, with the automatic detection and removal of long-range LD regions.

**Figure S3:**
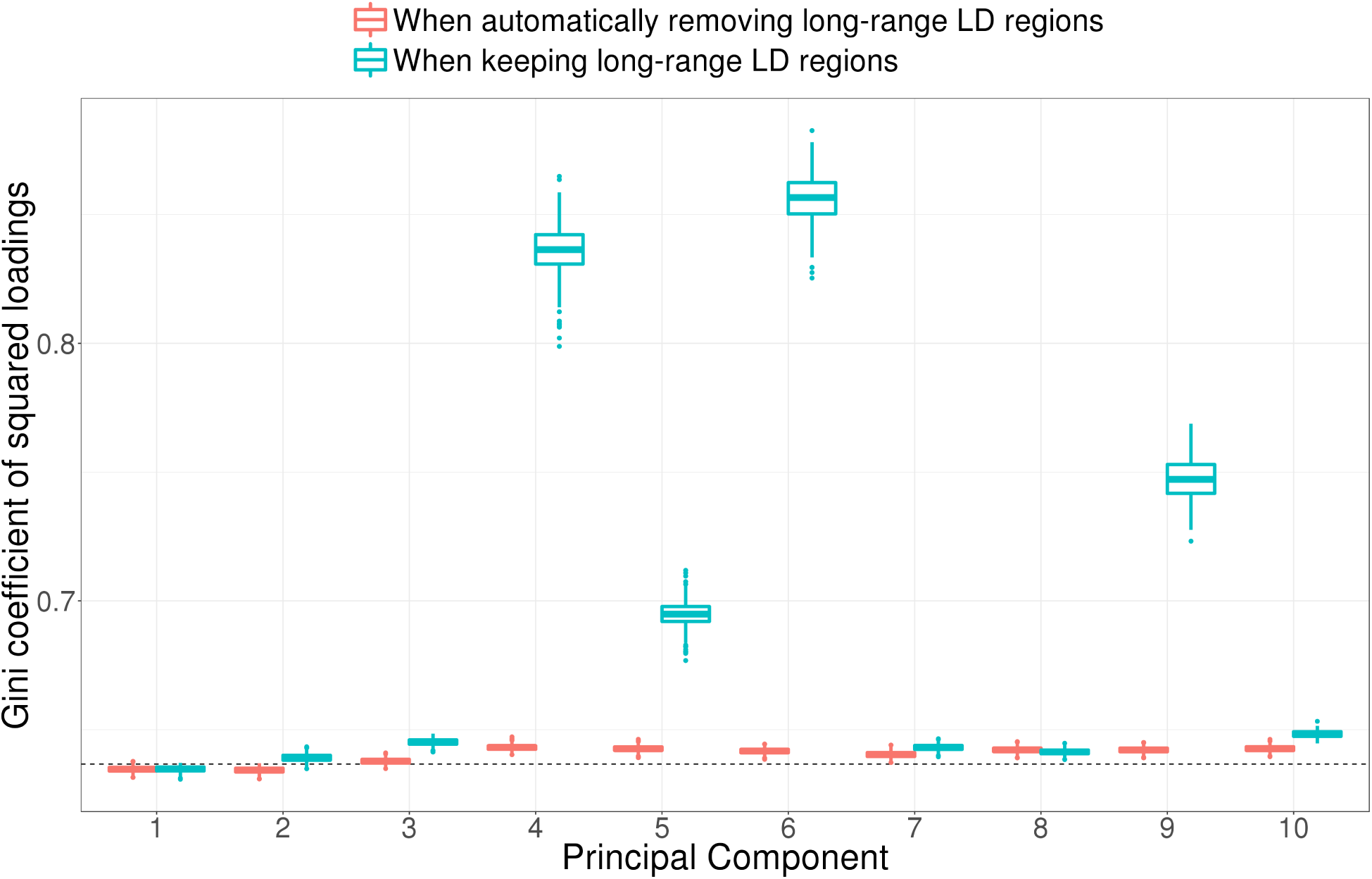
Boxplots of 1000 bootstrapped Gini coefficients (measure of statistical dispersion) of squared loadings without removing any long range LD region (only clumping at *R*^2^ > 0.2) and with the automatic detection and removal of long-range LD regions. The dashed line corresponds to the theoretical value for gaussian loadings.

**Figure S4:**
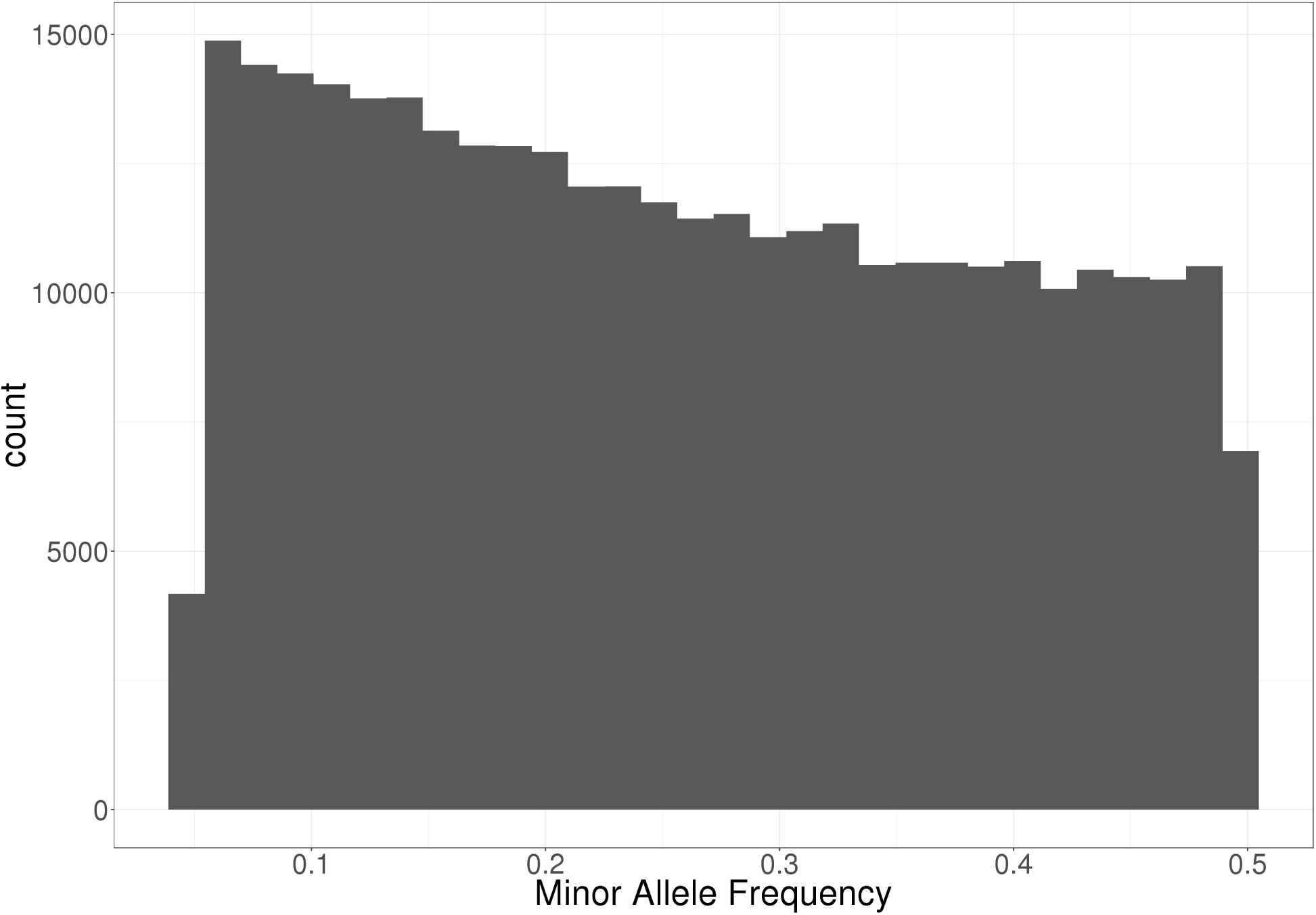
Histogram of the minor allele frequencies of the POPRES dataset used for comparing imputation methods.

**Figure S5:**
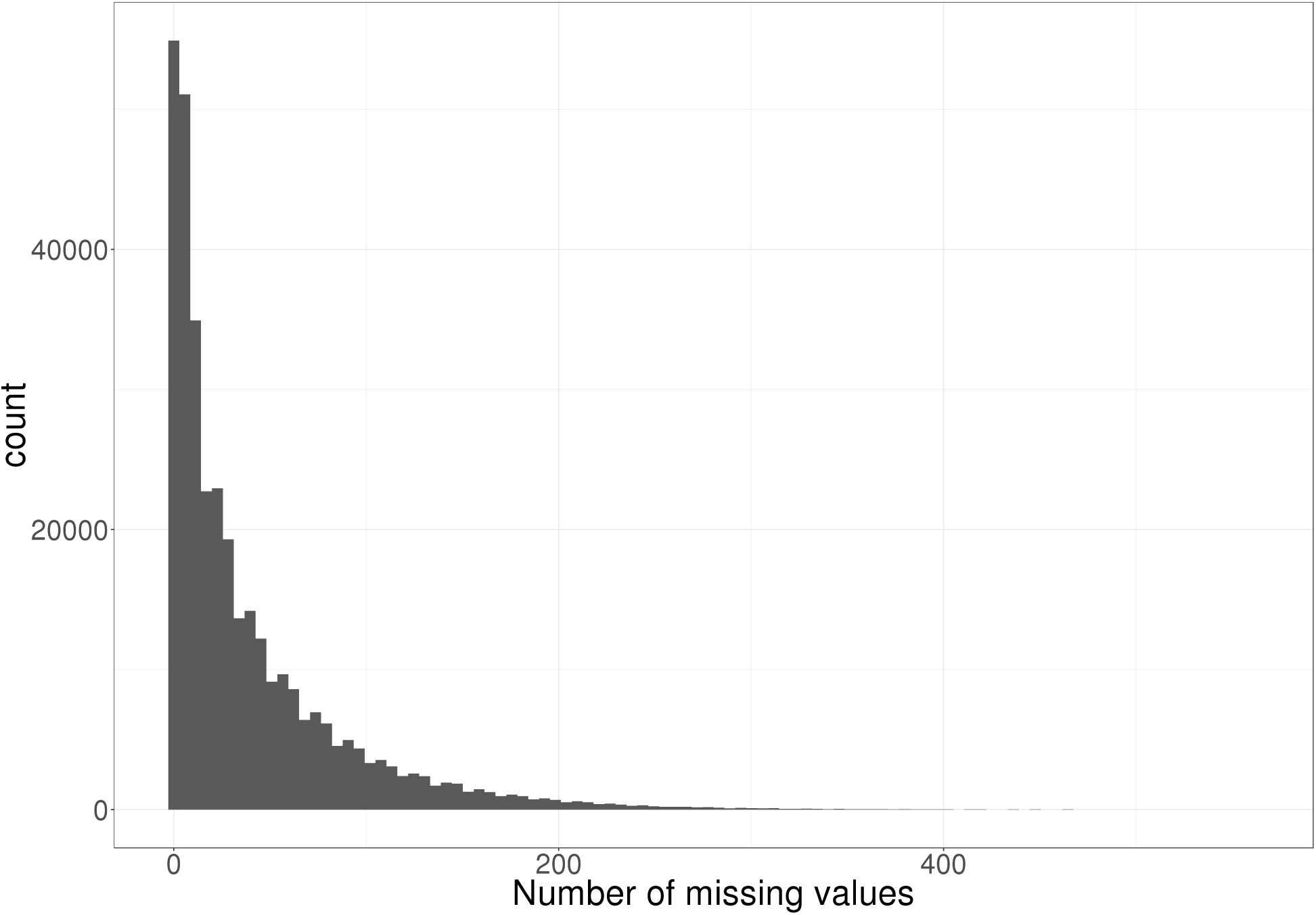
Histogram of the number of missing values by SNP. These numbers were generated using a Beta-binomial distribution.

**Figure S6:**
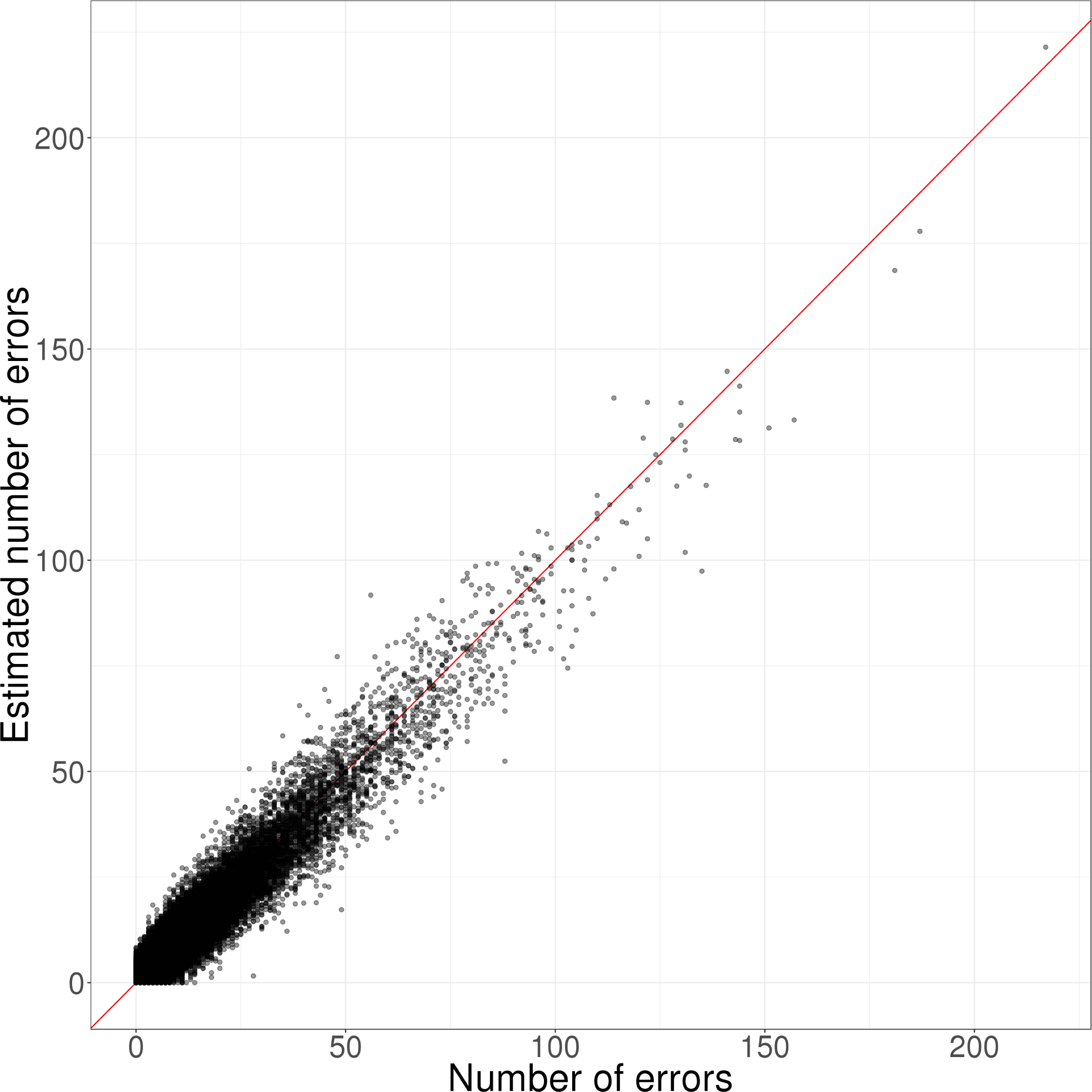
Number of imputation errors vs the estimated number of imputation errors by SNP. For each SNP with missing data, the number of imputation errors corresponds to the number of individuals for which imputation is incorrect. The estimated number of errors is a quantity that is returned when imputing with snp_fastimpute, which is based on XGBoost (Chen and Guestrin 2016).

## Footnotes

1 http://www.boost.org/

2 https://github.com/QuantGen/BEDMatrix

3 https://goo.gl/T5SJqM

4 https://goo.gl/8TngVE

5 https://goo.gl/8TngVE

